# Visual features contribute differently to preferences for different item categories

**DOI:** 10.1101/360172

**Authors:** Shiran Oren, Tal Sela, Dino J. Levy, Tom Schonberg

## Abstract

Low-level visual features have been known to play a role in value-based decision-making. However, thus far, mainly single features were tested on one type of item using one method of measurement. Here, we test the contribution of low-level visual features on three items types: fractal-art images, faces, and snack food items. We test the role of visual features on preferences using both subjective ratings and choices. We show that low-level visual features contribute to value-based decision-making even after controlling for higher level configural features of faces like eye-distance and market features of snacks like calories. Importantly, we show that while low-level visual features consistently contribute to value-based decision-making, different features contribute to different types of items when using different measurement methods. Our study highlights the necessity of using multiple item types and multiple measurement methods to construct a unifying framework regarding the contribution of low-level features to value-based decision-making.

## Visual features contribute differently to preferences for different item categories

Every day people make value-based decisions between different items. In this process, one needs to construct a representation of each item, assign a value to each item and choose the preferred item (Rangel, Camerer, & Montague, 2008). In the present study, we study to what extent do visual features of the options at hand influence this process? Understanding the role of low-level visual features in value computations is crucial to a variety of fields, from marketing to public health, as this understanding may help influence people’s preferences using subtle changes in low-level visual features.

When we observe an item, we putatively dissect it to its low-level characteristics, such as color (Livingstone & Hubel, 1988) and spatial frequency (De Valois, Albrecht, & Thorell, 1982; De Valois, Cottaris, Mahon, Elfar, & Wilson, 2000). Then, using the combination of such visual features, higher-level representations of the objects are formed and we identify the object (Riesenhuber & Poggio, 1999).

Several studies show the contribution of visual features to preferences of simple visual items. Color preferences were examined using color patches and found a general effect of higher preferences towards more saturated and cooler colors (i.e. blue preferred over red) both in a choice task (Hurlbert & Ling, 2007; McManus, Jones, & Cottrell, 1981) and in a scale rating task. A recent review of visual aesthetic preferences emphasized the roles of colors, spatial structures, and individual differences (Palmer, Schloss, & Sammartino, 2013). The influence of spatial frequency, which was shown to have a critical role in early visual processing (De Valois, Albrecht, & Thorell, 1982; De Valois, Cottaris, Mahon, Elfar, & Wilson, 2000) remains inconclusive regarding its contribution to preferences (Palmer et al., 2013). In another study that examined the contribution of visual features to preferences of natural and man-made scenes, scale-rating preferences were higher for images with higher quality (manifested as images with higher saturation, contrast & sharpness, and having less noise and grain) (Tinio & Leder, 2009). For images of snacks, items with higher luminance (Milosavljevic, Navalpakkam, Koch, & Rangel, 2012) or higher saliency (Towal, Mormann, & Koch, 2013) were more likely to be chosen in forced choice tasks. Although these studies showed effects for basic visual features on preferences for common objects, they each examined one visual feature on one type of object and thus, it is difficult to depict a clear picture of the relations between basic visual features and preferences for different types of objects.

For complex items such as faces, the configuration between facial features has been suggested to be critical in face processing (see Maurer, Le Grand, & Mondloch, 2002 for a review). Facial configuration was shown to play a role in face preferences (Cunningham, Roberts, Barbee, Druen et al., 1995; Geldart, Maurer, & Henderson, 1999). The width to height ratio (*fWHR*) (Weston, Friday, & Liò, 2007), was originally proposed as an evolutionary sexual marker (being greater for males) and was associated with aggressive and unethical behavior (e.g. Geniole, Keyes, Carré, & McCormick, 2014 but see Kosinski, 2017). Two other studies showed that changing the distance between local face elements (between the two eyes and between the eyes and nose) had a meaningful effect on preferences (Pallett, Link, & Lee, 2010). The distance between the eyes was also positively correlated with preference ratings (Cunningham, 1986).

A critical role for higher level features was also shown for food items, where, for example, the knowledge of a wine’s price (Plassmann, O’Doherty, Shiv, & Rangel, 2008) and of beer ingredients (Lee, Frederick, & Ariely, 2006) changed the items’ taste pleasantness ratings and neural activations. It has been shown that flavors associated with higher calorie foods induces higher preferences in adults, measured using a scale rating (Booth, Mather, & Fuller, 1982) and induced both higher preference ratings and actual consumption in children (Johnson, McPhee, & Birch, 1991). Thus, preferences for faces and foods are influenced by higher-level features as well as low-level features.

Still, it remains unknown if the effect of low-level visual features is the same as it was found for simple abstract stimuli such as patches of color or fractals. Moreover, it is not known if low-level visual features would have any effect on preferences, after controlling for the contribution of higher-level features. Despite ample research, the interplay between low-level visual features, item category, and methods of preference elicitation is unclear and has only been discussed in reviews or meta-analyses. It was not directly assessed, since most studies focused on a single visual feature, a single stimulus category and a single measurement method. Therefore, in the current research, we aimed to address this gap by systematically testing the influence of several low-level visual features of color attributes (*Hue*, *Saturation* and *Color*-*value*) and *Spatial*-*frequency*, on preferences for three categories of items. We used images of fractal-art, faces, and food snacks, as stimuli that contain different levels of higher-order complexity. Importantly, we examine the contribution of low-level features to preferences while adjusting for other non-visual attributes that play a role in preference formation. For fractal-art, we assume there are no higher attributes that affect preferences and examined only low-level visual features. For faces, we added three configural features (*fWHR*, *Eyes*-*distance* and *Nose*-*eyes distance*) and for snacks, we added three market features (*Calories*, *Product*-*weight*, and *Price*). Lastly, we examined preferences using both preference ratings and binary choices.

Thus, we aimed to address two main research questions: 1) what is the contribution of low-level visual features to preferences (measured by preference ratings) adjusted for higher-level features of different stimuli types; 2) how visual properties influence choices in a binary choice task, adjusted for the item’s preference ratings. Importantly, for robustness and generalizability, we used a large sample size collected across multiple studies in the laboratory and online, and report findings after a successful pre-registered replication of the online samples. We share all of our data as well as analyses codes.

## Methods

### Participants

A total of 1,036 participants took part in the experiment (see table 1 for sample details). All participants gave their informed consent in accordance with Tel Aviv University ethics committee and were paid for their participation.

**Table 1.** Demographic and sample details.

|  | Sample | Category | Age (sd) | % Females | n |
| --- | --- | --- | --- | --- | --- |
| Lab | (1) | Fractals | 24.38 (3.645) | 67% | 235 |
|  | (2) | Snacks | 24.64 (3.768) | 66% | 307 |
| Online | (3) | Fractals | 27.97 (5.388) | 51% | 107 |
|  | (4) | Snacks | 40.04 (13.112) | 50% | 108 |
|  | (5) | Faces | 42.47 (15.837) | 51% | 119 |
| Replications | (6) | Fractals | 38.41 (13.244) | 54% | 114 |
|  | (7) | Snacks | 39.04 (13.295) | 53% | 109 |
|  | (8) | Faces | 38.04 (12.818) | 45% | 115 |

### Stimuli

We used items from three different categories: 1) Sixty fractal artwork images obtained from the internet (“Fantastic Fractals”, 2013); 2) Sixty Israeli food snacks, all available in stores in Israel and cost up to 10 NIS (equal to ~$2.7) and 3) Sixty faces (30 females) obtained from the sibling database (Vieira, Bottino & Laurentini, 2013).

### Procedure

#### Laboratory experiments

(samples 1-2): We obtained preference ratings for fractals and snacks from 12 behavioral experiments (overall 342 participants) conducted in the lab (of which 8 experiments had both fractals and snacks ratings (n=200), one with only fractals (n=35) and 3 with only snacks ratings (n=107). All these experiments had a standard rating procedure, before any exposure to the rated items occurred. Given the rating task was similar across these 12 behavioral experiments, we examined the interaction between samples and the effects of interests, and found none of them to be significant. Thus, we combined all preference ratings in the laboratory as if they were collected from a large joint sample, one for fractals (sample 1) and the other for snacks (sample 2) (see table 1 for details). In the lab experiments, participants rated their preferences for the fractals using a continuous numerical scale, in which they indicated how much they liked each fractal from 1-10 (See Figure 1a). For the snack foods, participants indicated their willingness to pay (WTP) for each food item using the incentive-compatible Becker-DeGroot-Marschak auction (Becker, DeGroot, & Marschak, 1964), with a continuous price scale of 1-10 NIS. We used the WTP task as it is considered a task that elicits the participant’s preference ratings for the food snacks. Participants used a mouse to indicate their preference on a continuous number scale. The procedure was self-paced, and each item was presented once, resulting in 60 trials per participant, for each category. Faces were not rated in lab experiments. There were no binary choices in the lab experiments.

**Fig.1.**
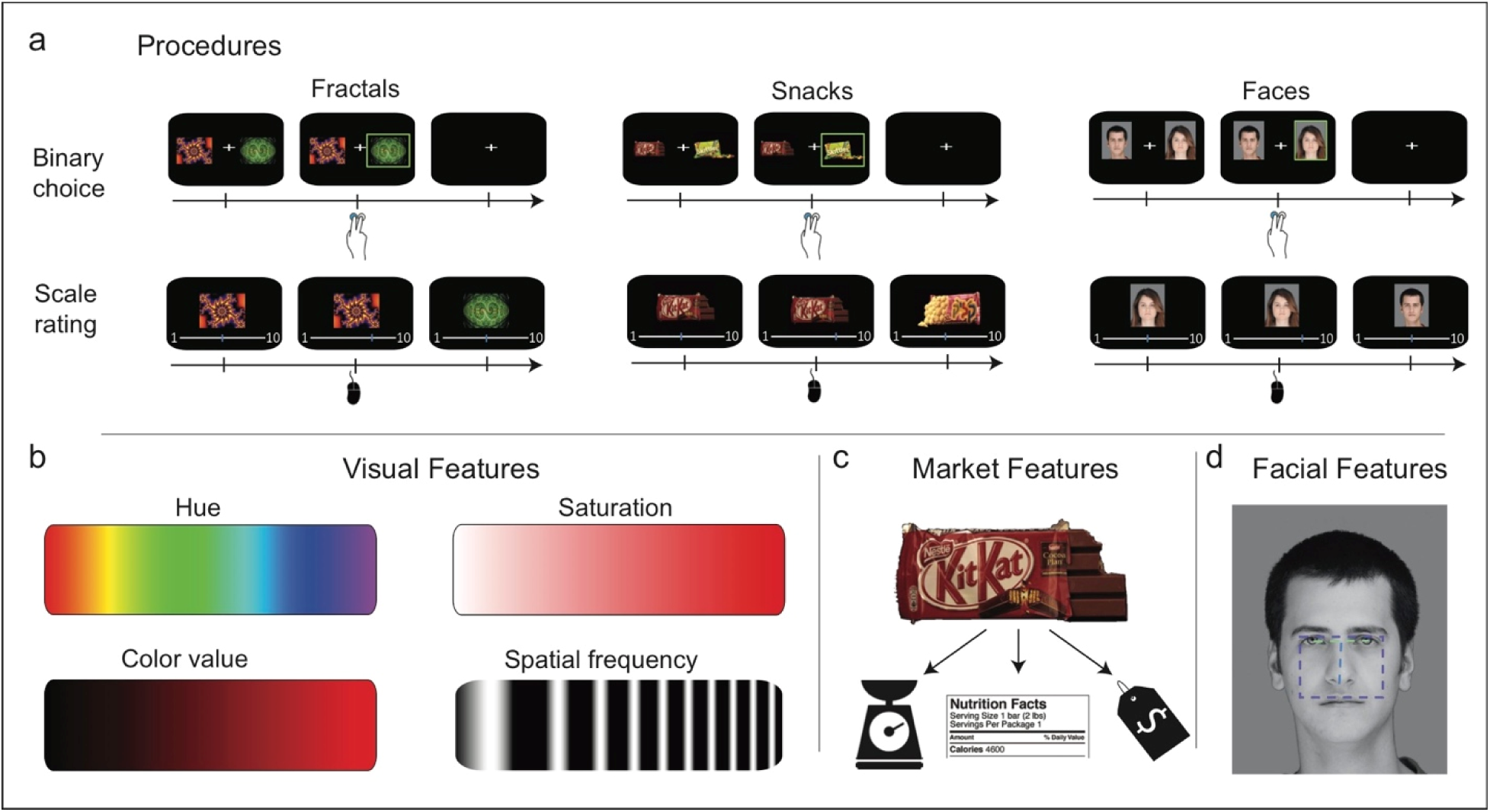
Study design. (a) Trial timeline in each of the two measurement methods and for the three different item types used. (b) The four low-level visual features extracted for all item types, ranging from lowest (left) to highest (right) value of the feature. (c) The three market features extracted for the food items. (d) The three facial features extracted for faces - *fWHR* is indicated by the purple rectangle’s aspect ratio, *Eyes*-*distance* by the green line and *Nose*-*eyes distance* by the blue line.

#### Online experiments

(samples 3-5): to replicate the lab preference ratings and explore the effects of visual features on binary choice tasks, we obtained choices and preference ratings in three online experiments (for fractals, snacks and faces). Overall, 334 participants took part in the online experiments, of which 107, 108 & 119 participated in the fractals, snacks and faces experiment, respectively. All online experiments were conducted via an Israeli online website (https://www.midgampanel.com/) that specializes in conducting online experiments. Each participant performed the binary choice task (See Figure 1b) followed by the preference rating procedure (See Figure 1b) of one of the categories (fractals, snacks or faces). In the binary choice task, ~14% of all possible binary choice combinations (60^∗^59/2, resulting in 240 trials per participant) were randomly selected and presented for each participant. On each trial, participants indicated which of the two items they preferred by pressing the keyboard. Each choice was presented for 2.5 seconds, followed by a 1 second fixation. The preference rating for all categories (fractals, snacks and faces) was obtained via the non-incentive scale rating procedure, identical to the lab preference rating procedure, described above in 2.1.

#### Replication experiments

(samples 6-8): to obtain a full replication of our data, we preregistered our data acquired from samples 1-5 and performed an identical replication of the above online experiments (https://osf.io/zmp49/?view_only=8c65101a29f14140b771aa87bcd91106). Overall, 338 participants took part in the replication online experiments, of which 114, 109 & 115 participated in the fractals, snacks and faces experiments, respectively.

### Feature analyses

*Visual* features: We extracted four visual features for each item using Matlab (Mathworks, Inc. Natick, MA, USA): *Hue*, *Saturation*, *Color*-*value* (color attributes according to the HSV color-map, calculated as the mean attribute of the item’s image (Joblove & Greenberg, 1978)) and *Spatial*-*frequency* (calculated as the mean image gradient using the Sobel-Feldman operator (Sobel & Feldman, 1968)).

In addition, we acquired the following category specific features for faces and snacks:

*Facial* features: We extracted three facial features for faces using the Viola-Jones algorithm (Viola & Jones, 2001) using Matlab: 1) *Eyes*-*distance* (the distance between the two eyes normalized by face size); 2) Facial width to height ratio (*fWHR*); 3) *Nose*-*eyes distance* (the distance between the bottom of the nose and the center of the two eyes, normalized by face size). *Market* features: We collected three market features for each of the snacks: 1) *Price*; 2) *Product*-*weight* (in grams); 3) *Calories* (per 100g). We extracted the information from the labeling on the snack’s package and from the internet.

### Behavioral analyses

Both the WTP scores and the preference ratings were z-scored separately for each participant, to remove variance between participants resulting from them using different ranges of the scale. All the extracted values of the features (*visual*, *market* and facial features) were also z-scored to enable a direct comparison of regression coefficients. We removed from further analysis items with feature values exceeding 3 standard deviations (SDs) away from the mean (1 fractal, 2 snacks, and 1 face). We excluded trials with reaction times exceeding 3 SDs away from the mean (calculated within task and within participant) or trials with no response. Overall, we removed an average of 2.31% (0.349 SDs) of trials per participant, across all samples. In addition, we removed participants with more than 30% excluded trials, in either rating or choices data. We removed 16 participants in the online and replication samples (2-4 participants for each sample), concluding overall 1014 valid participants in all samples. No participants were removed from the lab samples. We performed all analyses in R (version 3.3.2).

### 1: Is there a linear relationship for low-level visual features and category-specific features, with preference ratings?

To examine the influence of low-level visual features and category-specific features (market and facial features) on preference ratings, we fitted for each category a linear mixed-effects regression model (model 1, see supplementary material), with a random-intercept and random slopes. That is, we allowed the intercept and the slope coefficients of each of the features (visual and category-specific features) to vary across participants. Ratings served as the dependent variable and the different features as fixed and random independent variables. We fitted this model separately for each of the lab (samples 1-2), online (samples 3-5) and replication (samples 6-8) samples. For each of the samples, we entered all features together to the regression model. Thus, the results reflect the unique contribution of each feature adjusted for all other features.

### 2: Do low-level visual features and category-specific features affect binary choices, adjusted for preference ratings?

To examine the influence of the different features on choices, adjusted for preference ratings, we first determined the preference ratings of each item for each participant using their ratings in the scale rating task. We then calculated the value difference between the two items in every choice option (hereafter *delta value* (e.g., Milosavljevic, Navalpakkam, Koch, & Rangel, 2012)). We chose to account for value differences in each choice option, since they are expected to have a strong influence on choice. By entering *delta value* into the regression models, we allowed the exploration of more subtle effects of visual features on choices. Furthermore, for each of the features (visual and category-specific features) we extracted the score difference between the two items (left item minus right item) in every choice option (hereafter *delta feature*, e.g. *delta Hue*, *delta Price* etc.). For each of the samples, we entered the delta value feature with all delta visual and delta category-specific features together to the regression model. We fitted for each category a mixed-effects logistic regression (model 2, see supplementary material). We fitted a random-intercept and random-slope model, with choices as the dependent variable, and *delta value* and all *delta features* (the delta of visual, market and facial features) as fixed independent variables. We allowed for the intercept and slope of *delta value* to vary across participants. We fitted this model separately for each of the online (samples 3-5) and replication (samples 6-8) samples.

Data and code sharing: All data and analyses codes are available here: https://osf.io/zmp49/?view_only=8c65101a29f14140b771aa87bcd91106

## Results

The current study is composed of many samples with numerous possible effects, and thus we report below the summary of effects. Detailed description of all model results with effect sizes and confidence intervals are reported in the supplementary material.

### 1: Is there a linear relationship for low-level visual features and category-specific features, with preference ratings?

In ratings data obtained from the experiments conducted in the lab, we found that each of the visual features had a different influence on preference ratings, and in some cases an opposite effect, depending on the items’ category (Figure 2a). Specifically, *Hue* had a positive effect on preference ratings of fractals, whilst a negative effect on snacks. *Saturation* had a negative effect only for snacks and *Color*-*value* had a negative effect only for fractals. *Spatial*-*frequency*, however, had a positive effect for both fractals and snacks.

**Fig.2.**
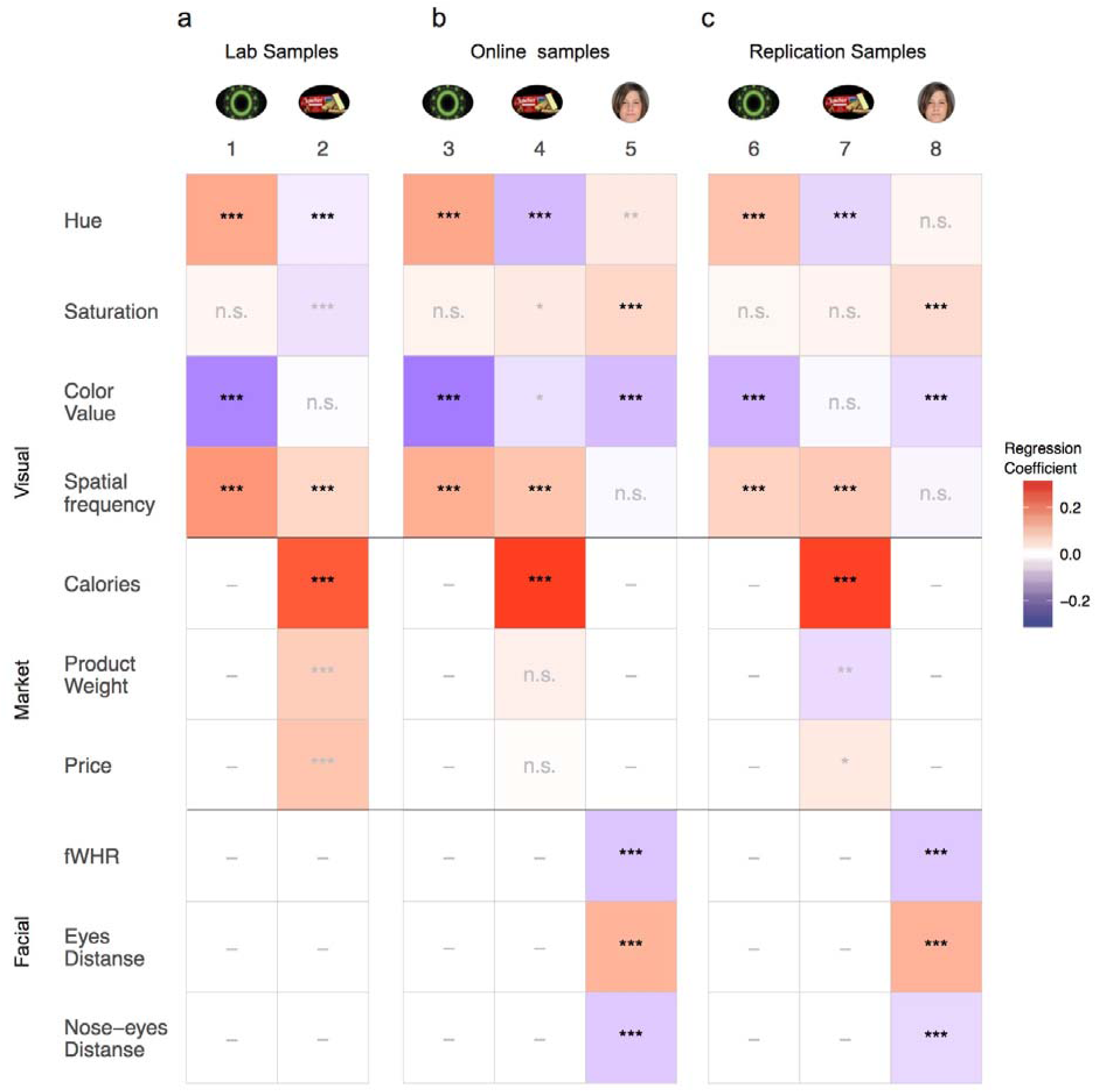
Results summary of the mixed linear regression models for ratings of lab (a), online (b) and replication (c) samples. Each column represents different samples and each row represents a different feature. The color of the square indicates the coefficient value for the current feature in the current samples’ model, from high (red) to low (blue). Black text indicates that the current feature was significant across all samples in the category. Gray text indicates that one or more samples of this category were not significant, thus the effect was not stable across all samples. ^∗^ p<0.05; ^∗∗^ p<0.01; ^∗∗∗^ p<0.001.

In order to examine the robustness of these results we repeated the lab experiments on an online cohort of participants. This complex pattern of relations between visual features and item category in their influence on preference ratings was mostly replicated in the online experiments (Figure 2b). That is, *Hue* had a positive effect on fractals *(sample 3)* and a negative effect on snacks *(sample 4). Color*-*value* had a negative effect on fractals *(sample 3)* and *Spatial*-*frequency* had a positive effect on fractals and on snacks *(samples 3 and 4).* The negative effect for *Saturation* on snacks in the lab data was not replicated and even reversed to a positive effect in the online samples. In addition, we found a negative effect for *Color*-*value* on snacks, which was absent in the lab samples.

In addition to the samples of fractals and snacks, in the online experiments we collected data for preference ratings for faces. Similar to the fractals and snacks, we found that there was an effect of visual features on preference ratings for faces. However, in general these were different features and the direction of influence was different compared to the effect we found for fractals and snacks. That is, *Saturation* and *Hue* had a positive effect and *Color*-*value* had a negative effect on preference ratings for faces (*sample 5*). These results further support our finding that the effect of visual features on preference ratings is category specific.

We then examined the replication results of our pre-registered samples, which serve as a third replication of the effects of visual features on preference ratings for fractals and snacks (following the lab and online samples), and a second replication for faces (following the online sample). Importantly, we replicated the complex pattern we obtained in the lab and online samples (Figure 2c). Specifically, *Hue* had a positive effect on fractals *(sample 6*, *similar to samples 1 & 3)* and a negative effect on snacks *(sample 7*, *similar in samples 2 & 4). Color*-*value* had a negative effect on fractals *(sample 6*, *similar in samples 1 & 3)*, and *Spatial*-*frequency* had a positive effect on fractals and on snacks *(samples 6 and 7*, *similar to samples 1 and 2 & 3 and 4).* For faces, *Saturation* had a positive effect and *Color*-*value* had a negative effect (*sample 8*, *similar to sample 5*). The negative effect for *Saturation* on snacks in the lab data *(sample 2)* was not replicated in the replication samples *(sample 7*, *similar to sample 4).* The effects for *Saturation and Color*-*value* on snacks *(sample 4)*, and for *Hue* on faces *(sample 5)* that were found in the online samples, were not replicated in the replication samples *(samples 7 and 8)*.

In order to investigate the role of basic visual features adjusted for the effects of higher level features, we added market features for snacks and facial features for faces to the regression models, alongside the visual features. The effects of the higher-level features were examined and replicated to some extent. For snacks, we found a positive effect for *Calories*, *Product*-*weight* and *Price* in the lab sample *(sample 2).* The effect of *Calories* on ratings was replicated first in the online sample *(sample 4)* and again in the replication sample after pre-registration *(sample 7).* However, the effects of *Product*-*weight* and *Price* were not replicated in the online sample *(sample 4).* In the replication sample *(sample 7)*, the effect of *Price* was replicated, but the effect of *Product*-*weight* was reversed. For faces, we found a negative effect for *fWHR*, a negative effect for *Eyes*-*distance* and a positive effect for *Nose*-*eyes distance* in the online sample *(sample 5).* These effects for faces were fully replicated in the replication sample after pre-registration *(sample 8)*.

Overall, the category-dependent pattern for the influence of visual features on preference ratings was stable across independent samples and experimental settings (lab vs. online experiments). For detailed results of the regression models see table s1 supplementary material.

### 2: Do low-level visual features and category-specific features affect binary choices, adjusted for preference ratings?

We next examined the influence of low-level visual features and category-specific features on actual choices after we controlled for their value, as indicated in their preference ratings. Similarly, to the preference ratings results, the effect of the various features on choice was category specific. That is, each feature affected choice differently and this effect depended on the specific items’ category (Figure 3). Note, that these effects are after controling for the higher-level features in the model (facial and market), and were mostly replicated in the independent replication samples (samples 6, 7, and 8). It is important to emphasize, that the effects of the various features on choice (described in Figure 3), exist after adjusting for the item’s preference ratings, as the ratings of each item were included in the regression model. That is, visual features can impact choices, regardless of the items’ value. Particularly, participants tended to choose items with higher *Hue* in fractals *(sample 3)*, higher *Saturation* in faces *(sample 5)*, lower *Color*-*value* in snacks *(sample 4)*, higher *Spatial*-*frequency* in fractals *(sample 3)* and lower *Spatial*-*frequency* in faces *(sample 5).* For the category-specific features, participants tended to choose items with higher *Calories* and higher *Price* in snacks *(sample 4)*, higher *Eyes*-*distance* in faces *(sample 5)* and lower *Nose-eyes distance (sample 5).* All these effects were than replicated in the replication samples (*sample 6* for fractals, *7* for snacks and *8* for faces), except for the effect for *Nose*-*eyes distance*, which was not replicated *(sample 8).* In addition, we found a negative effect for *Product*-*weight* on snacks *(sample 7)*, which was not found in the online sample *(sample* 4). These results indicate a pattern, by which different visual features influence choices between items, in a replicated manner within the same category but not across categories. For detailed results of the regression models see table s2 supplementary material.

**Fig.3.**
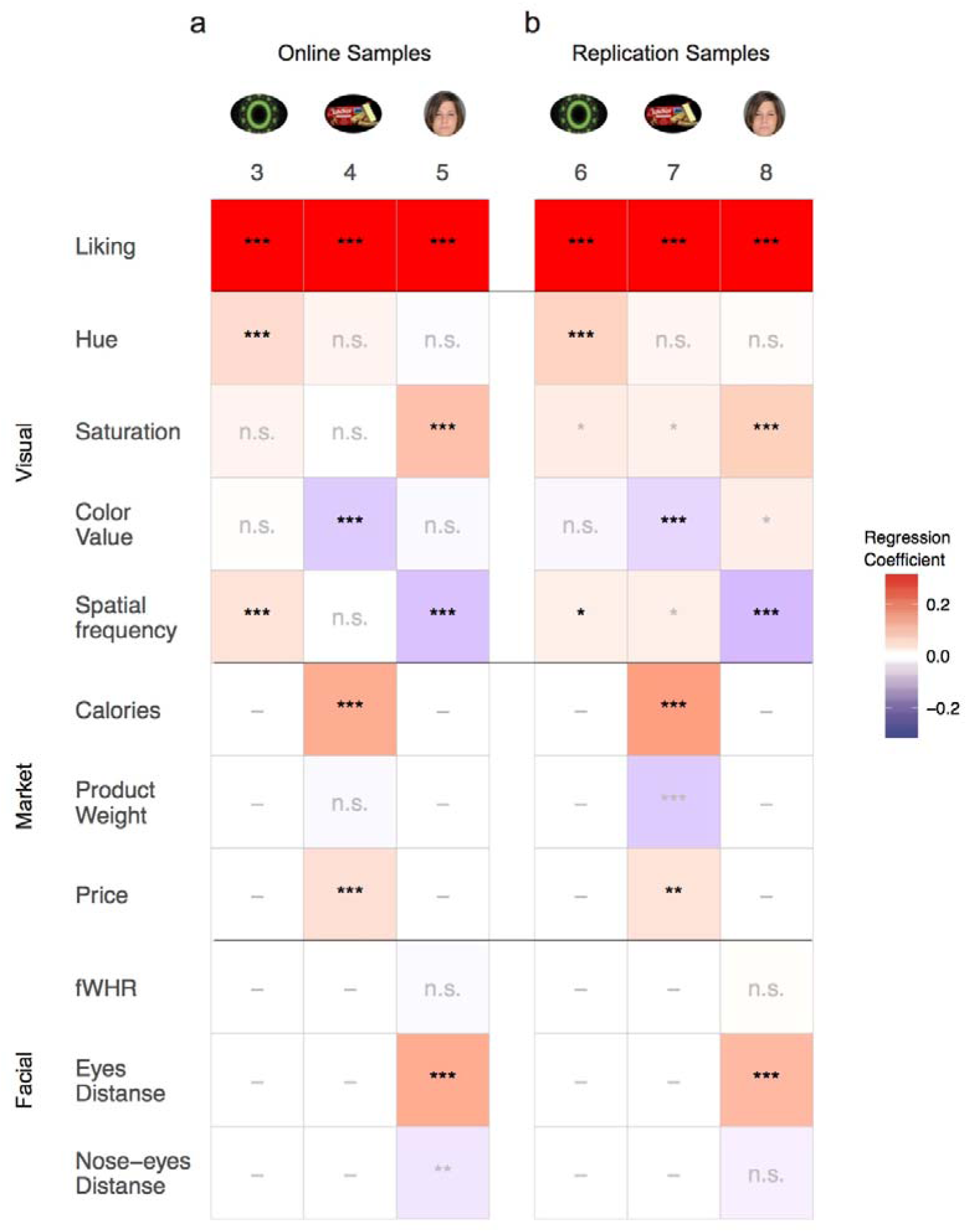
Results of the mixed logistic regression models for choices for online (a) and replication (b) samples. Each column represents different samples and each row represent different features. The color of the square indicates the coefficient value for the current feature in the current sample’s model, from high (red) to low (blue). Black text indicates that the current feature was significant across all samples of this category. Gray text indicates that one or more samples of this category were not significant, thus the effect was not stable across samples. p<0.05; ^∗∗^ p<0.01; ^∗∗∗^ p<0.001.

### Effects across tasks

We demonstrate a complex pattern of the effects of basic visual features on preference ratings and binary choices. Figure 4 shows a summary of the effects that were replicated, obtained only in preference rating (4a), only in choices (4b), and in both procedures (4c). There were several effects that were similar across both task procedures. For the basic visual features, *Hue* and *Spatial*-*frequency* had a positive effect on fractals, and *Saturation* had a positive effect on faces. For the higher-level features, *Eyes*-*distance* (Facial) had a positive effect on faces, while *Calories* (Market) had a positive effect on snacks. Hence, these effects are stable across independent samples (including a replication of pre-registered samples) and could be generalized across measurement procedures.

**Fig.4.**
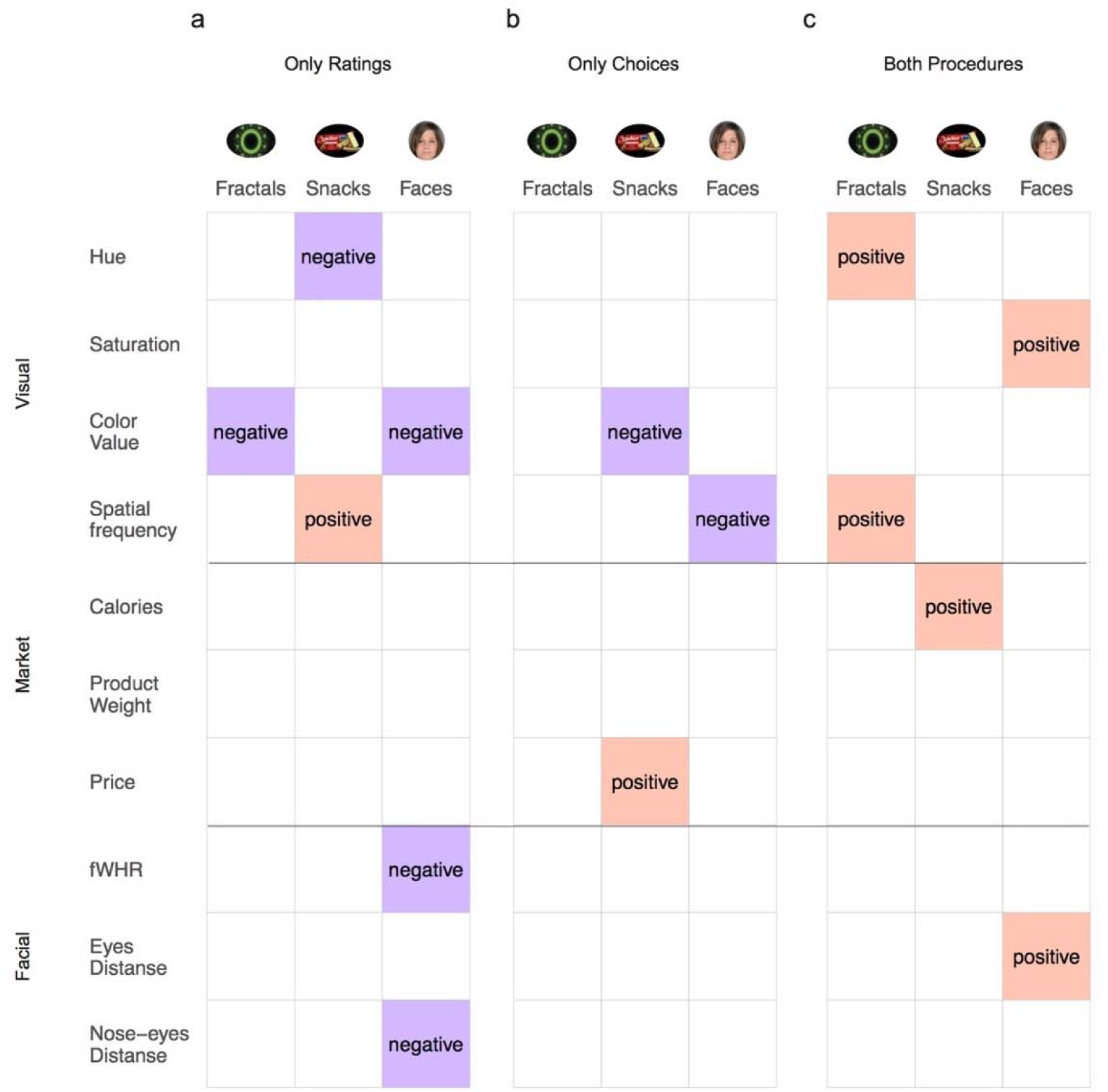
Summary of results from both ratings and choice data. All effects shown here were stable across all samples for each category, for only ratings (a), only choices (b) or both ratings and choices (c). Columns represent categories and rows represent features. The color indicates the direction of the effect (red: positive, blue: negative).

In contrast, there were several effects that were replicated only in one task procedure, but not in the other. In the preference ratings task, *Color*-*value* had a negative effect for fractals and faces, *Hue* had a negative effect on snacks, and *Spatial*-*frequency* had a positive effect on snacks. For *the higher*-*level* features, only facial features replicated in the preference ratings task, as *fWHR* and *Nose*-*eyes distance* had a negative effect on preference ratings.

On the other hand, in the choice task, *Spatial*-*frequency* had a negative effect on faces, while *Color*-*value* had a negative effect for snacks. Note, that there was no clear and replicated effect on fractals in the choice task that was absent in the ratings task. For *the higher*-*level* features, only market features replicated solely in the choice task, as *Price* had a positive effect on choices.

These results further emphasize the uniqueness of the effects, by which different visual features influence preferences or choices between items, in a replicated manner within the same category but not across categories or across measurement procedures. We note that when testing the role of visual features on binary choice we accounted for the subjective ratings of each item. Therefore, the influence of a feature on ratings is statistically accounted for when we examine the effect of that feature on binary choices. Meaning, the differences between measurement tools implies that the visual features influences each measurement differently.

## Discussion

In the current study, we test for the first time in one study, the influence of low-level visual features on preferences of fractal art images, faces and snack-food items, using both ratings and binary choices. We show that low-level visual features contribute to preferences, but differently for each stimulus type and measurement method. Most studies thus far showed the influence of isolated visual properties, such as contrast or hue, on specific aesthetic items (such as paintings, abstract images, etc. (Palmer et al., 2013). However, an investigation of isolated visual properties does not provide the possibility to examine the contribution and interactions of many different features in parallel and different measurement methods.

We focused on four basic visual features, which have a critical role in low level visual processing: the three main color features – *Hue*, *Saturation* and *Color*-*value* (Livingstone & Hubel, 1988), and *Spatial*-*frequency* (De Valois, Albrecht, & Thorell, 1982; De Valois, Cottaris, Mahon, Elfar, & Wilson, 2000). Most studies that examined color preferences on simple color patches stimuli had shown a general preference toward cooler and brighter colors (i.e. higher hue, *Saturation* and *Color*-*value* (Hurlbert & Ling, 2007; McManus, Jones, & Cottrell, 1981; Palmer & Schloss, 2010). However, we show that for complex items, this tendency was replicated only for the *Hue* of fractals and *Saturation* of faces. Namely, participants preferred fractals with higher *Hue* (but not faces or snacks) and faces with higher *Saturation* (but not fractals or snacks), in both rankings and choices. Moreover, we found the opposite effect for *Color*-*value*, which showed a negative effect on rankings of fractals and faces, and on choices of snacks. Interestingly, we also found an opposite effect for Hue on ratings of snacks, showing higher preferences for lower hues. Based on the current literature it is difficult to find definitive reasons for why a specific feature contributed to one category over the other. This is one of the main conclusions of our study: when testing only a certain feature using a certain methodology on a specific stimulus type (as was done in previous studies) it is hard to generalize across domains and features. For example, the *Spatial*-*frequency* of an image, despite having an important role in visual processing, has not been examined thoroughly with relation to its effect on visual preferences (Palmer et al., 2013). In the current study, we show that *Spatial*-*frequency* indeed has a stable influence on visual preferences, but in a category-specific manner. That is, *Spatial*-*frequency* had a positive influence on both ratings and choices of fractals, whereas it had a positive effect on ratings of snacks and a negative effect on choices of faces.

Furthermore, our results provide interesting insights regarding the effects that higher-level features have on preferences and choice. For snacks, we found that participants like items with higher *Calories*, as indicated both in their ratings and choices, in line with other studies that showed higher preferences for high-calorie foods (Booth et al., 1982; Johnson et al., 1991). In addition, choices were influenced by the snacks’ *Price*, in accordance with studies showing the effect of price expectations on preferences (Lee et al., 2006; Uher, Treasure, Heining, Brammer, & Campbell, 2006). Still, the lack of effect of *Price* on preference rating is interesting and calls for further investigation. For faces, participants preferred faces with greater distance between the eyes and nose in both ratings and choices, in line with previous work (Cunningham, 1986). Additionally, participants preferred faces with lower nose to eyes distance and smaller *fWHR*, but these two features had no effect on binary choices above and beyond ratings. The effects of these two features were inconclusive in previous studies. The nose to eyes distance was shown to correlate with preferences in an inverted U-shape (Pallett et al., 2010), whilst the Midface range (corresponding with this feature) had no correlation with preference (Cunningham, 1986). For *fWHR*, a recent large sample study claimed for a null effect for this feature (Kosinski, 2017). Thus, our study sheds light on the interaction between facial features and preferences, however, further studies are required to determine the exact role of these features on preferences.

Following the replication crisis in different scientific fields (Lithgow, Driscoll, & Phillips, 2017; Nave, Camerer, & McCullough, 2015; Prinz, Schlange, & Asadullah, 2011), the research community has become committed to producing reproducible science, using larger sample sizes, pre-registrations, sharing open codes and data (Borsboom, Green, & Mabry, 2015; Nosek, Spies, & Motyl, 2012). Therefore, here we share all our data, analysis and task codes. Furthermore, the findings of our samples have been pre-registered prior to collecting additional data and we were able to replicate their findings in three new samples. This provides further robustness and generalizability of our results.

To conclude, we offer for the first time an elaborate testing of multiple visual features on multiple categories with several measurement tools to show that the influence of low level visual features is complex, and specific to the item category tested and the way we estimate items’ value (either by preference ratings or choice). Moreover, we demonstrated that low-level features affect preference ratings and also influence choices even after controlling for preference ratings, showing that these effects are sustainable and independent of the items’ value. Importantly, we were able to replicate almost all effects of the low-level visual features that we found in the current study, demonstrating that these effects are stable and could be generalized across samples. This exemplifies the importance of pre-registration and testing our results using independent samples in order to obtain robust conclusions. Future studies are suggested to follow this approach of testing multiple features, on multiple items in different settings potentially on the same participants, to take into account as many features as possible to be able to shed light on the long-lasting question posed by Fechner (1871) regarding the influence of visual features on preferences.

Author Contributions
S.O. designed experiments, collected and analyzed data and wrote the manuscript. T.Se. assisted with design of the experiments, data analysis and contributed to write-up of manuscript. D.L. contributed to the design of the experiments, data analysis and manuscript write-up; T.Sc. designed experiments, assisted with data analysis and wrote the manuscript with S.O. All writers actively contributed to the writing process of the manuscript.

## Acknowledgements

This work was supported by the Israeli Science Foundation (ISF number 1798/15 and 2004/15) and the European Research Council (ERC) under the European Union’s Horizon 2020 research and innovation program (grant agreement n° 715016) granted to Tom Schonberg. We would like to thank Jeanette Mumford for statistics advice.

